# AC-BioSD : A biomolecular signal differentiator module with enhanced performance (extended version)

**DOI:** 10.1101/2024.01.29.577841

**Authors:** Emmanouil Alexis, José L. Avalos, Luca Cardelli, Antonis Papachristodoulou

## Abstract

Temporal gradient estimation is a pervasive phenomenon in natural biological systems and holds great promise for synthetic counterparts with broad-reaching applications. Here, we advance the concept of *BioSD* (*Biomolecular Signal Differentiators*) by introducing a novel biomolecular topology, termed *Autocatalytic-BioSD* or *AC-BioSD*. Its structure allows for insensitivity to input signal changes and high precision in terms of signal differentiation, even when operating far from nominal conditions. Concurrently, disruptive high-frequency signal components are effectively attenuated. In addition, the usefulness of our topology in biological regulation is highlighted via a PID (Proportional-Integral-Derivative) bio-control scheme with *set point weighting* and filtered derivative action in both the deterministic and stochastic domains.

## I. INTRODUCTION

Estimating time derivatives of biomolecular signals is an essential process of several natural biological systems. Bacterial chemotaxis [1] stands as a notable example. This navigation mechanism guides bacteria, such as *Escherichia coli*, towards beneficial chemicals (*attractants*) and away from harmful ones (*repellents*). It relies on a temporal sampling strategy enabling bacteria to calculate temporal gradients in order to infer spatial ones.

Designing reliable biomolecular topologies capable of (mathematically) differentiating signals, such as the concentration of biochemical substances or light, can be pivotal in addressing challenges in the realm of Synthetic Biology [2]–[4]. For instance, such topologies can be utilized as speed biosensors to monitor the rate of change with respect to biomolecular signals of interest. These biosensors are therefore able to assess cellular uptake and secretion rates of biomolecules, rendering them powerful tools for bioremediation purposes [5], [6] or for establishing efficient interactions within synthetic microbial communities for a variety of applications [7]–[10]. In parallel, derivative action allows the development of advanced regulation strategies. For example, PID (Proportional-Integral-Derivative) control constitutes one of the cornerstones of control engineering [11]. Recent *in-silico* studies have compellingly showcased the benefits of derivative control within PID bio-control schemes, such as enhanced dynamic performance and mitigated stochastic noise. These schemes include implementations based on nucleic acid strand displacement systems, single cells as well as microbial consortia [12]–[18].

A convenient and effective way of realizing derivative action in a biological setting is by leveraging the concept of *BioSD* (*Biomolecular Signal Differentiators*), initially introduced in [19]. *BioSD* motifs can serve as tunable, general-purpose signal differentiator modules around their nominal operation. Under appropriate tuning, they are able to compute the first derivative of a biomolecular signal with high accuracy. At the same time, they offer protection against undesired amplification of high-frequency signal components through an embedded second-order low-pass filter. Additionally, they provide considerable experimental flexibility, and their output is encoded as a species concentration which simplifies measurement procedures. *BioSD* modules can also be employed for derivative control, potentially in conjunction with proportional and integral action (PID control) [17]. Note that different approaches to realizing derivative action (including derivative control) via biochemical interactions have been proposed in the recent literature [12]–[16], [18], [20]–[22]. For a concise comparative discussion, we refer the reader to [23].

In this work, we expand upon the *BioSD* concept by introducing a novel biomolecular topology with enhanced capabilities, termed *Autocatalytic-BioSD* or *AC-BioSD*, using the *BioSD-III* topology [19] as its structural foundation. The distinguishing feature of *AC-BioSD* lies in the fact that the input excitation is applied through an autocatalytic biochemical reaction. Notably, this modification does not add structural complexity in terms of the number of reactions and species. However, it endows *AC-BioSD* with significant advantages. Firstly, contrary to *BioSD-III*, the ability of the former to differentiate (or attenuate) input signals is independent of the input excitation reaction, allowing fine-tuning based exclusively on its intrinsic characteristics. At the same time, its behavior away from the nominal operation (non-local behavior) remains remarkably reliable, whereas that of *BioSD-III* can be greatly compromised. Furthermore, we demonstrate how *AC-BioSD* can be used for derivative control by adapting the PID scheme described in [17] and make comparisons with PI control as well as with the same PID scheme based on *BioSD-III*. Finally, it is important to emphasize that this study considers only realistic (bounded) physical signals of finite time duration which can be considered both differentiable and Fourier-transformable [19], [24].

The paper is structured as follows: Section II provides a brief background encompassing modeling principles, computational approaches, and the concepts of *BioSD* action as well as a special PID bio-control strategy. Section III presents the structure and the local behavior of the *AC-BioSD* module. Section IV examines its non-local ability to differentiate signals and Section V concentrates on derivative control. Lastly, Section VI summarizes our work and discusses future experimental and theoretical research avenues.

## II. BACKGROUND

### A. Modeling principles and computational analysis

We use three main types of biochemical reactions to structurally describe the biomolecular topologies herein, which are aligned with the law of mass action [19], [25], [26]: general transformation of reactants into products, catalytic production and catalytic inhibition. Assuming *A, B* are two biochemical species, these reactions take the following general form *A →B, A → A* + *B, A* + *B → A*, respectively. The corresponding graphical representations are given in Fig. 1(a). Note that autocatalytic production can be regarded as a special case of catalytic production where a reactant enables its own formation, potentially in the presence of some other reactant with or without the latter being consumed - for example *A* + *B → A* + 2*B* or *A* + *B →* 2*B*, respectively.

**Fig. 1:**
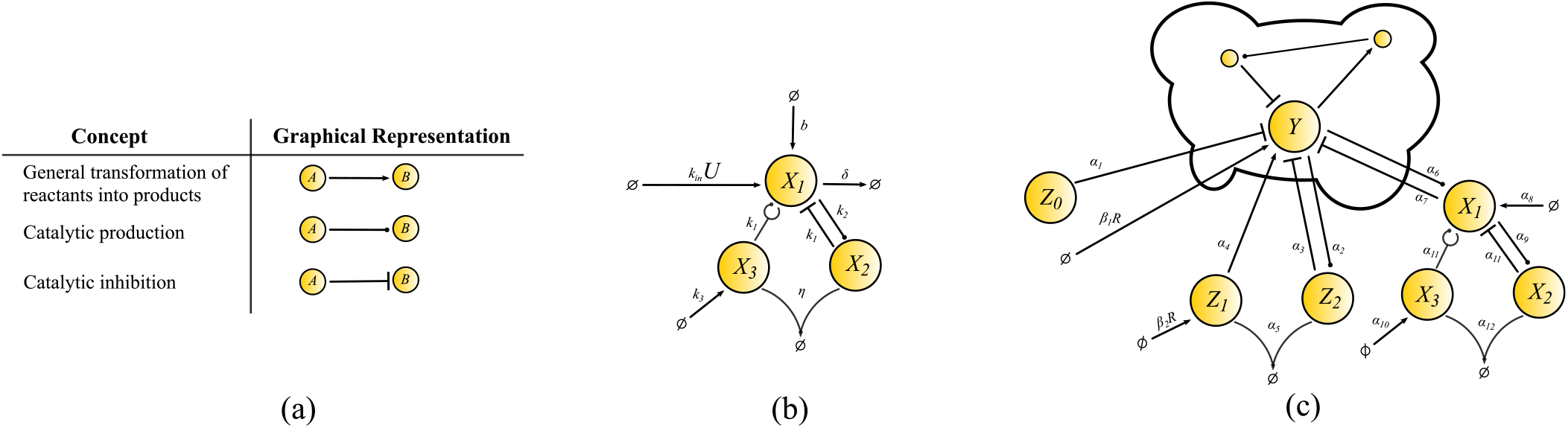
(a) Biochemical reactions considered in this work [25]. (b) *BioSD-III* module with *U* and *X*_1_ being the input and the output signal/species, respectively [19]. (c) PID control architecture with *set point weighting* and filtered derivative action [17]. The cloud network represents an arbitrary biomolecular process including the target/output species *Y* . The controller species are shown outside the cloud network.

To study the deterministic dynamics of the topologies (Sections III, IV, V), we use Ordinary Differential Equation (ODE) models and perform simulations in MATLAB (Mathworks). In Section V we also perform simulations in the chemical reaction simulator Kaemika [27] where stochasticity is supported through the Linear Noise Approximation (LNA) of the Chemical Master Equation (CME) [28].

### B. BioSD-III module

The architecture of *BioSD-III* [19] is illustrated in Fig. 1(b) and its dynamics can be modelled as:

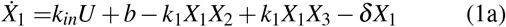

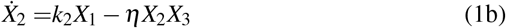

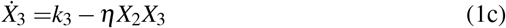

where *k*_*in*_, *b, k*_1_, *k*_2_, *k*_3_, *δ, η ∈* ℝ_+_. Here *U* is the input excitation process, corresponding to one of the following reactions: *U → X*_1_ or *U → U* + *X*_1_ (at rate *k*_*in*_); these can be summarized by the notation 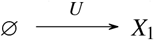 as in Fig. 1(a).

We now discuss some important characteristics of *BioSD-III* which are proved in [19]. Given any non-negative constant input *U*^*\**^, system (1) has a unique positive steady state 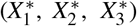 which is locally exponentially stable. Assuming sufficiently small deviations *u, x*_1_, *x*_2_, *x*_3_ around 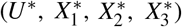, its input/output relation can be described in the Laplace domain by the transfer function:

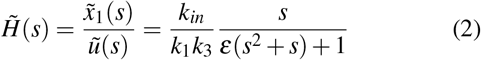

where *s* is the Laplace variable (complex frequency) and 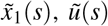 are the Laplace transform of *x*_1_, *u*, respectively. Moreover:

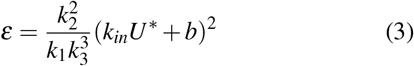

Equation (2) represents an ideal signal differentiator (accompanied by a constant gain) in series with a second-order low pass filter. Moving now to the frequency response of *BioSD-III*, we obtain via Fourier Transform:

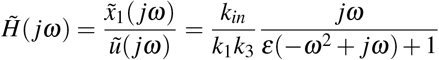

where *ω* is the (real) frequency, 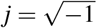 is the imaginary unit and 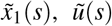 are the Fourier transform of *x*_1_, *u*, respectively.

Finally, in the time domain, for a sufficiently slow input signal of sufficiently long time duration, the output of *BioSD-III* can be approximated by:

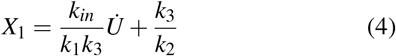

### C. PID control and BioSD action

Fig. 1(c) depicts a biochemical reaction network implementation of a PID control scheme with *set point weighting* and filtered derivative action, as introduced in [17]. Here, derivative control is carried out by *BioSD-III* (see species *X*_1_, *X*_2_, *X*_3_), which interacts with the output species, *Y*, via a production-inhibition loop. *BioSD-III* contributes to the proportional control involved, as well.

The corresponding closed-loop dynamics is given by the following set of ODEs:

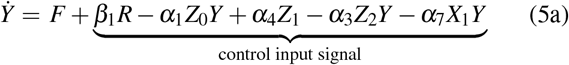

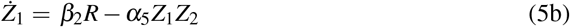

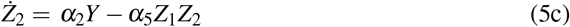

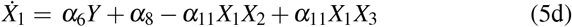

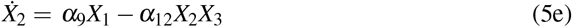

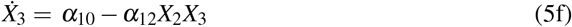

where *α*_*i*_ ∈ ℝ_+_ with *i* ∈ ℕ and 1 ≤ *i* ≤ 12. *R* is a non-negative reference signal, *β*_1_, *β*_2_ are non-negative scaling parameters and *Z*_0_ is an auxiliary species with constant concentration. *F* denotes potential interactions with respect to *Y* inside the cloud network. Furthermore, for simplicity, the “self-degradation” rate of *X*_1_ (see term *− δ X*_1_ in Equation (1a)) is assumed to be zero here as it is generally not necessary for the proper function of the *BioSD* modules (the same is true for *AC-BioSD* introduced in the next section) [17], [19].

Given some constant value of the reference signal, *R*^*\**^, we assume the existence of a (locally) asymptotically stable and biologically meaningful steady state (indicated by ^*\**^ next to the respective variables) of system (5). Focusing on the closed-loop behavior around this steady state, we further assume sufficiently small deviations of the (uppercase) variables which are represented by the corresponding lowercase letters. As shown in [17], the control input signal in Equation (5a) can take the following form of a PID control law with *set point weighting* and filtered derivative action:

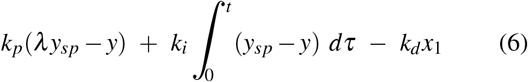

where: 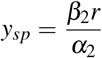 (set point), 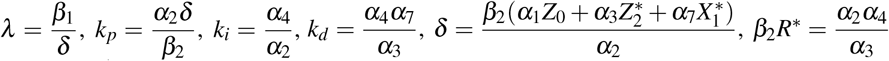

Furthermore:

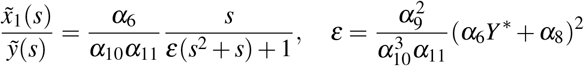

## III. *AC-BioSD* module

Fig. 2(a) illustrates the biomolecular architecture of *AC-BioSD*. It consists of the same biochemical reactions as *BioSD-III* except for the part of the input excitation process. Here, the latter corresponds to one of the following autocatalytic reactions: 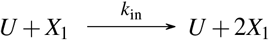 or 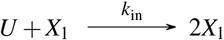. Note that the function of *AC-BioSD* remains the same regardless of which of the above reactions is adopted.

**Fig. 2:**
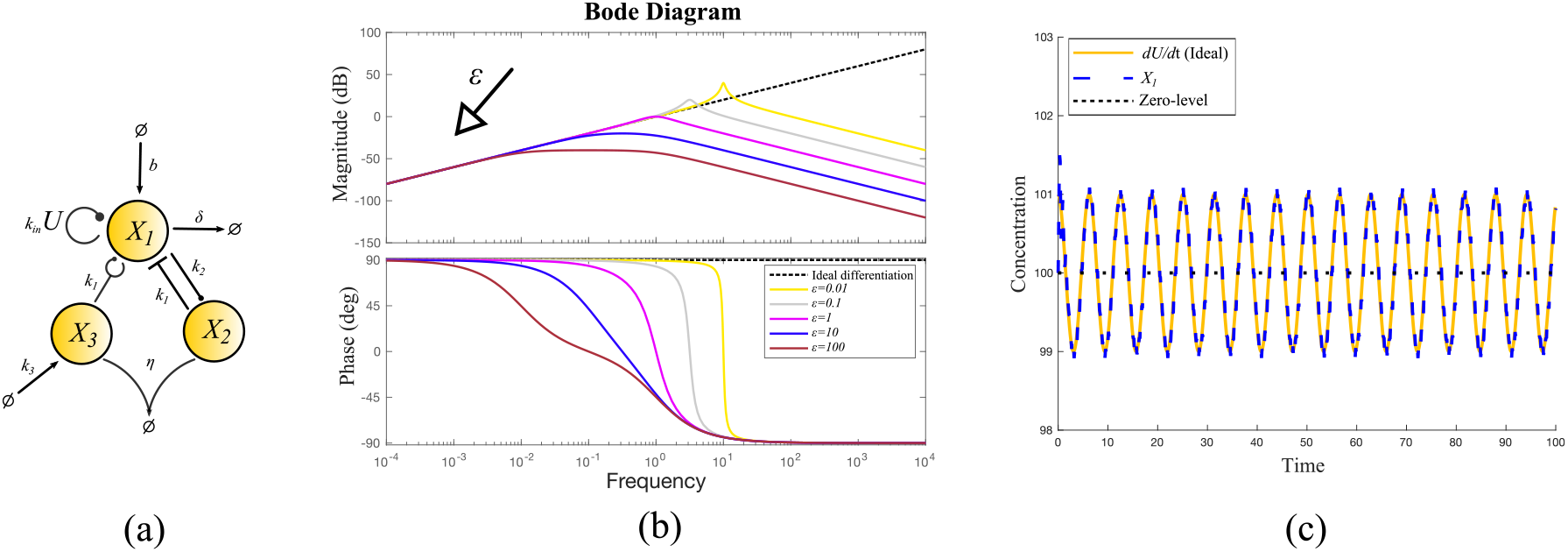
(a) *AC-BioSD* module with *U* and *X*_1_ being the input and the output signal/species, respectively. (b) Bode plot based on Equations (15), (16). For simplicity we assume 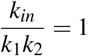. (c) Simulated output response of *AC-BioSD* (system (7)) with *k*_*in*_ = 1, *b* = 220, *k*_1_ = 1, *k*_2_ = 1, *k*_3_ = 100, *η* = 30. Equation (11) therefore gives *ε* = 0.0484. Moreover, we apply the input signal *U* +*U*_*h f n*_ for 0 ≤ *t* ≤ 100, where *U* = 1.2 + *sin*(*ωt*) represents the useful information (signal to be differentiated) and *U*_*h f n*_ = 0.2*sin*(400*ωt*) represents unwanted high-frequency noise (signal to be attenuated). Note that the output response is practically the same to the one of the *BioSD* modules in [19] (see Fig. 5.A) where the same *ε* and input signal are considered.

The dynamics of *AC-BioSD* is captured by the ODE model:

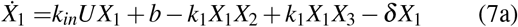

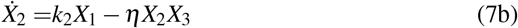

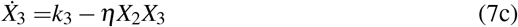

where *k*_*in*_, *b, k*_1_, *k*_2_, *k*_3_, *δ, η* ∈ ℝ_+_.

### Theorem 1

For any positive constant input *U*^*\**^, system (7) has a unique positive steady state which is locally exponentially stable.

*Proof:* By setting the time derivatives of system (7) to zero and taking into account that *X*_1_, *X*_2_, *X*_3_ are always non-negative as physical quantities, we get the following unique steady state:

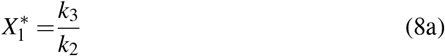

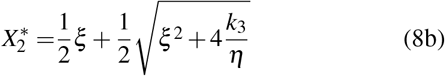

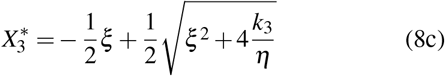

with

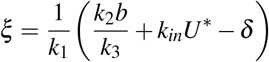

Note that *k*_*in*_, *b, k*_1_, *k*_2_, *k*_3_, *δ, η ∈* ℝ_+_ which also entails 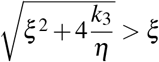. Clearly, the steady state (8) is positive.

Jacobian linearization of system (7) around the fixed point (8) yields:

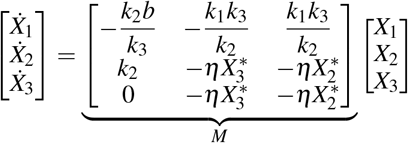

We calculate the characteristic polynomial of *M* as:

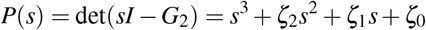

with 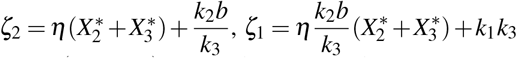 and 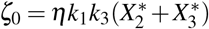. According to Routh-Hurwitz criterion, *P*(*s*) has all roots in the open left half-plane if and only if *ζ*_2_, *ζ*_1_, *ζ*_0_ *>* 0 and *ζ*_2_*ζ*_1_ *> ζ*_0_. The first condition clearly holds.

For the second one we have:

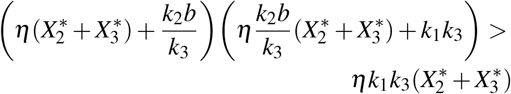

or

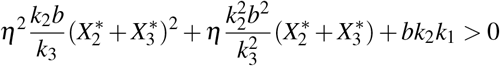

which is always true as a sum of positive quantities. Consequently, *M* is Hurwitz and steady state (8) is locally exponentially stable for system (7).

### Proposition 1

In the neighbourhood of 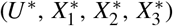, the input/output relation of system (7) in the Laplace domain is given by 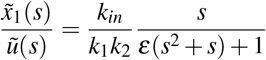 with 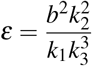.

*Proof:* We adopt the coordinate transformations: 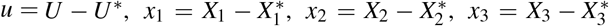 denoting sufficiently small deviations around 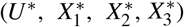. Subsequently, through Jacobian linearization around the aforementioned fixed point, the local dynamics of system (7) is derived as:

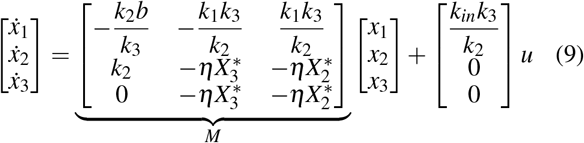

By adopting the linear transformation *ϕ* = *x*_2_ − *x*_3_, system (9) becomes:

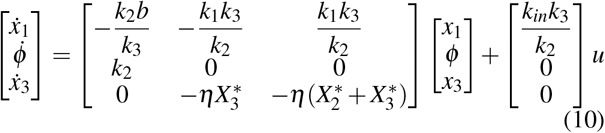

We next introduce the nondimensional variables: 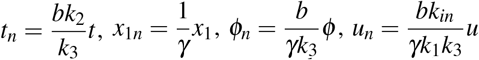, where *γ* is an arbitrary scaling parameter that carries the same units as *x*_1_. We also consider the nondimensional parameter

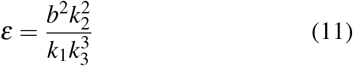

Hence, focusing on the dynamics of *x*_1_ and *ϕ* in system (10), we obtain: 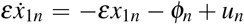 and 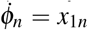 or, equivalently:

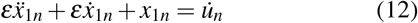

Applying Laplace Transform to Equation (12) results in the following transfer function:

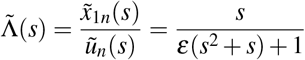

If we now take into account the aforementioned nondimensional variables, we get:

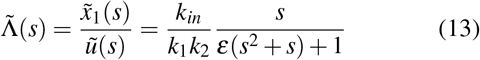

To study the frequency response of system (13) in the above proof, we apply Fourier Transform. Due to the existence of asymptotic stability, we can do that by setting *s* = *jω* to obtain:

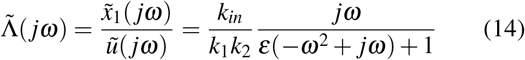

or, equivalently, 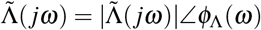 with:

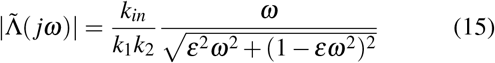

and

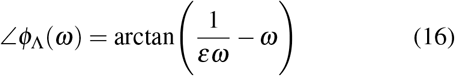

Perfect signal differentiation entails 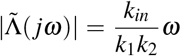 and ∠*ϕ*_Λ_(*ω*) = 90^*°*^ whereas perfect signal attenuation entails 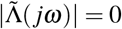.

Taking also into account the initial coordinate transformations as well as Equation (8a), the output of *AC-BioSD* in the time domain can be approximated as:

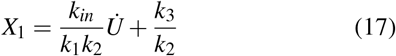

assuming a sufficiently slow input signal of sufficiently long time duration. Note that 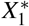 can be regarded as our “x-axis” or “zero-level concentration” around of which signal differentiation is carried out.

As can be seen in Fig. 2(b)-(c), for a given positive *ε*, there is a “low-frequency” and a “high-frequency” range over which signal differentiation and signal attenuation is effectively performed, respectively. As *ε* increases, the latter expand towards the former (and vice versa). Different performance standards can therefore be met by appropriately tuning this parameter.

It is apparent that both *BioSD-III* and *AC-BioSD* exhibit filtered derivative action and their behavior around their nominal operation can be described by equations of the same form but characterized by some distinct parametrization differences. That is, the reaction rates involved generally influence the behavior of these modules in different ways (Equation (4) vs (17)). As already pointed out, the nondimensional parameter *ε*, which is a function of several reaction rates, plays a pivotal role for the appropriate tuning and, by extension, the reliable operation of the *BioSD* modules. As Equations (3), (11) indicate, in the case of *BioSD-III, ε* highly depends on the input excitation process since it is a function of both *k*_*in*_ and *U*^*\**^. On the other hand, in the case of *AC-BioSD ε* is completely independent of that process. This particular attribute can be a major benefit for a circuit designer implementation-wise. More specifically, the topology of *AC-BioSD* can be designed to achieve certain performance requirements based solely of its internal reactions, eliminating the need of a priori knowledge of the input signal and/or the input reaction rate. This stands in contrast to *BioSD-III*, where the presence of different input signals/reaction rates may necessitate the re-tuning of other internal reactions of the topology (potentially during operation) which can be challenging in many biological settings [2]–[4].

## IV. Non-local signal differentiation

Under suitable tuning, *BioSD-III* can act as a signal differentiator of high accuracy in the vicinity of its equilibrium point [19]. Nevertheless, as the system is driven away from this point the accuracy might drop and, in sufficiently distant operating regimes, the ability of signal differentiation can be entirely compromised. This happens because, as the system moves further from its equilibrium point, its inherent non-linearities gradually become significant, changing its (non-local) behavior in an unfavorable manner. Interestingly, this is not the case for *AC-BioSD* whose signal differentiation capacity far from its equilibrium point shows remarkable consistency. In Fig. 3 and 4 we computationally demonstrate the above by applying appropriate input signals which allow us to investigate such operating regimes.

**Fig. 3:**
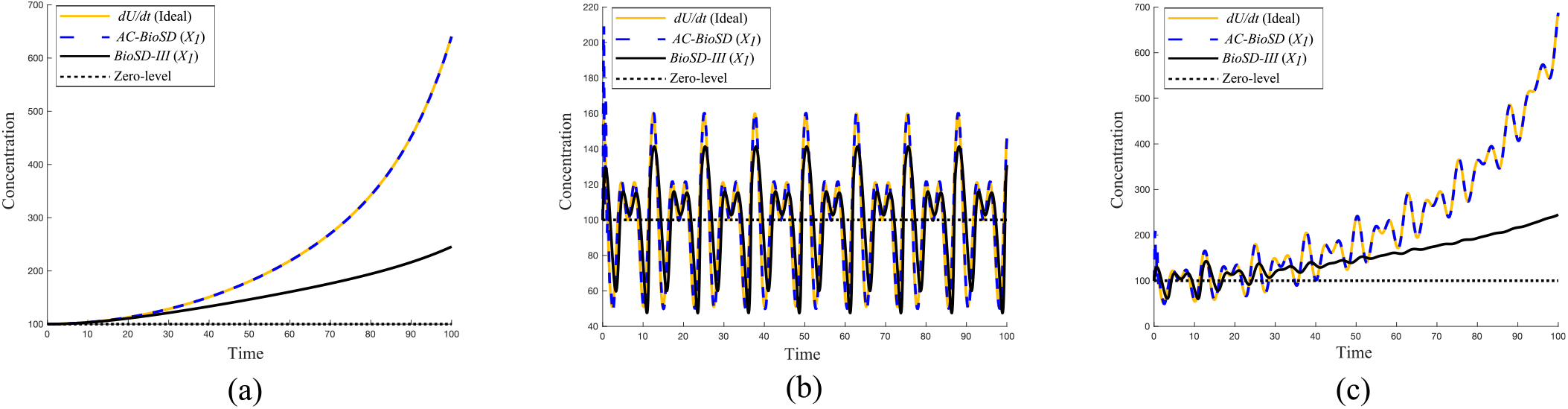
Simulated output response of *AC-BioSD* (system (7)) and *BioSD-III* (system (1)) to the following input signals: (a) *e*^0.08*t*^ + 0.01(*t*^3^ + *t*^2^ + *t* + 1) (b) 60 + 30*sin*(*t*) + 20*sin*(1.5*t*) (c) the sum of (a), (b) for 0 ≤ *t* ≤ 100. The paramater tuning corresponds to the performance standards adopted in Fig. 2(c) (for details regarding *BioSD-III* see Fig. S1.B and Fig. 5.A in [19]).

**Fig. 4:**
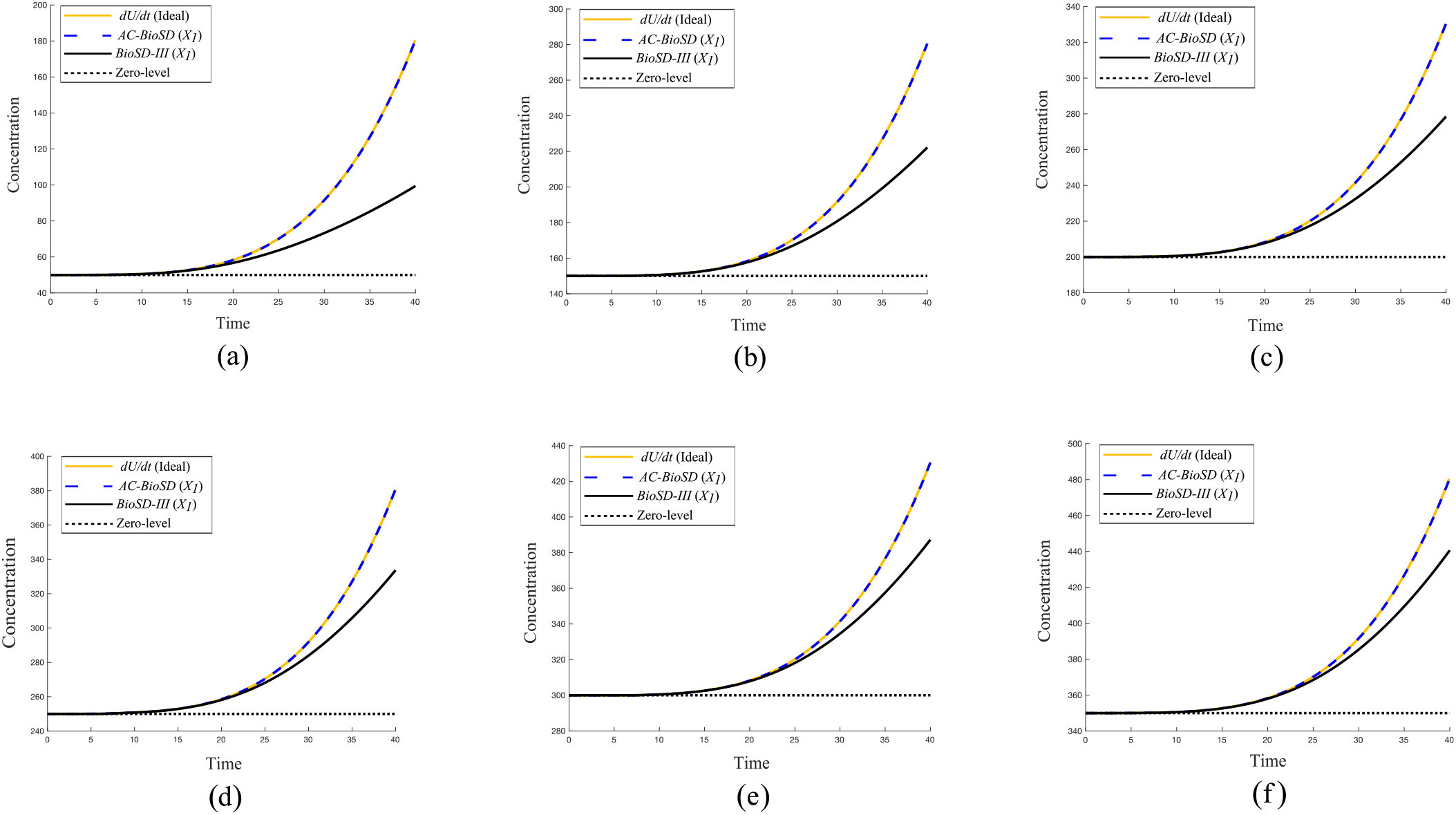
Simulated output response of *AC-BioSD* (system (7)) and *BioSD-III* (system (1)) to the input signal 0.00001(*t*^5^ + *t*^4^ + *t*^3^ + *t*^2^ + *t*) for (a) *ε* = 0.1 (b) *ε* = 0.05 (c) *ε* = 0.01 (d) *ε* = 0.005 (e) *ε* = 0.001 (f) *ε* = 0.0005. In all the depicted scenarios we have: *U*^*\**^ = 0, *k*_1_ = 1, *k*_2_ = 1,*η* = 30 while *b* and *k*_3_ vary appropriately. For simplicity, the output gains of *AC-BioSD* and *BioSD-III* are chosen to be equal to 1, i.e. 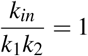 and 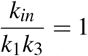, respectively. A shorter simulation time is used here compared to Fig. 3 to better illustrate the region around the point where the signal differentiation ability of *BioSD-III* begins to diminish. Note though that the high output accuracy of *AC-BioSD* remains uncompromised for longer time intervals.

## V. *AC-BioSD* derivative control

Fig. 5(a) illustrates how derivative control can be realized via the *AC-BioSD* module within the PID control scheme presented in Section II.C. For demonstration purposes, we consider a simple biological process of two mutually activated species [17], [25] inside the cloud network (this does not affect the generality of the PID architecture outside the cloud network). The corresponding closed-loop dynamics is given by the ODE model:

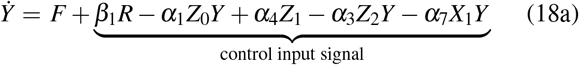

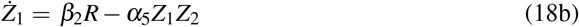

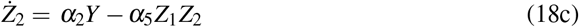

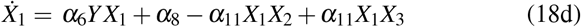

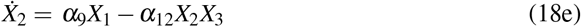

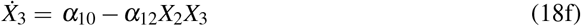

where *F* = *γ*_1_ *− γ*_3_*Y* + *γ*_6_*W*, 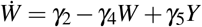 and *γ*_*m*_ ∈ ℝ_+_ with *m* ∈ ℕ and 1 ≤ *m* ≤ 6.

**Fig. 5:**
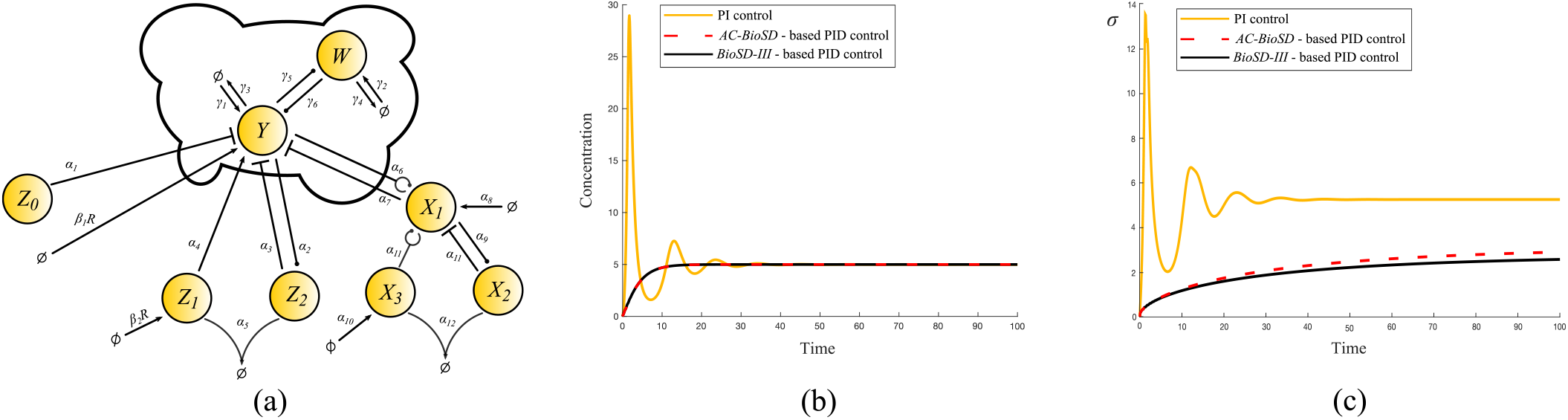
(a) Biochemical reaction representation of system (18). For comparison reasons, the simulated output responses and standard deviations depicted in (b) and (c), respectively, refer to the scenario considered in [17] (Fig. 4) using the same parameter values. Note that for the case of the *AC-BioSD*-based PID control (system (18)) we select *α*_6_ = 1 instead of *α*_6_ = 100 (which is used for the case of the *BioSD-III*-based PID control). This facilitates comparison as the respective derivative gains become equal.

Although, as expected, the dynamics of the derivative part of system (18) is different compared to the one in system (5), the remaining part of the two systems is identical. Consequently, using the methodology of Section III in [17], it is straightforward to show that the control input signal in Equation (18a) can be (locally) described by Equation (6) with the following modification regarding the derivative part (see Section III):

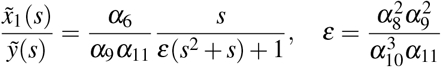

As illustrated in Fig. 5(b), both the *AC-BioSD*- and the *BioSD-III*-based PID controller practically yield the same output response in the deterministic setting, characterized by smooth transient dynamics - the overshoot and the oscillations observed in the PI control case are eliminated by the anticipatory derivative action. Simultaneously, Fig. 5(c) shows that both PID controllers, when compared to the PI controller, significantly reduce the standard deviation of the output, denoted here as *σ*, at steady state. However, in this specific example, the standard deviation in the *AC-BioSD* case is slightly elevated compared to the one in the *BioSD-III* case.

## VI. Conclusion

In this study, we present the *AC-BioSD* module, a novel *BioSD* design for signal differentiation in biomolecular environments. We built upon the architecture of the previously proposed *BioSD-III* module and appropriately embedded an autocatalytic input excitation process. Unlike the latter module, this structural modification renders the characteristics of *AC-BioSD*’s function insensitive to changes of the input reaction and offers greater reliability away from nominal conditions. Moreover, *AC-BioSD* can be conveniently employed for derivative control contributing to improved dynamic performance and suppression of stochastic noise.

We anticipate that the design approach introduced here will facilitate the experimental implementation of robust and predictable synthetic circuits. Notably, *AC-BioSD* consists of a finite set of (unimolecular and bimolecular) reactions following mass-action kinetics and, therefore, is suitable for molecular programming applications. For example, detailed information on how such a topology can be realized *in vitro* using strand displacement DNA-based chemistry is available in [12], [25]. In terms of *in vivo* implementations, the core parts of *AC-BioSD* (which are similar to the original *BioSD* modules) can be realized through gene expression systems – see, for instance, [19] for potential implementations in *Escherichia coli*. As for the autocatalytic reaction, it could be realized through a gene expression system based on positive feedback [26]. A promising candidate might be a protein, such as a transcription factor, which activates its own expression in the presence of a second chaperone protein [29] or light [30]. Potential limitations imposed by saturation nonlinearities in the models stemming from the aforementioned gene expression processes need to be taken into account, as well.

An intriguing theoretical extension of this work would involve exploring further the *AC-BioSD* concept using other *BioSD* modules, namely *BioSD-I* or *BioSD-II* [19], as structural foundations. Subsequently, investigation and comparison of different ways for applying derivative control, for instance within PID regulation strategies, using various *(AC- ) BioSD* modules would be of great interest. Performance characteristics of the output response that merit attention include deterministic features such as settling time, rising time, and overshoot as well as estimation of the variance or higher moments in the stochastic setting. Finally, our understanding of the non-local behavior of the *BioSD* modules would be enhanced by adopting more sophisticated mathematical approaches based, for example, on frequency-domain analysis methods for nonlinear systems [31], [32].

